# Agricultural and geographic factors shaped the North American 2015 highly pathogenic avian influenza H5N2 outbreak

**DOI:** 10.1101/645408

**Authors:** Joseph T. Hicks, Dong-Hun Lee, Venkata R. Duvuuri, Mia Kim Torchetti, David E Swayne, Justin Bahl

## Abstract

The 2014 – 2015 highly pathogenic avian influenza (HPAI) H5NX outbreak represents the largest and most expensive HPAI outbreak in the United States to date. Despite extensive traditional and molecular epidemiological studies, factors associated with the spread of HPAI among midwestern poultry premises remain unclear. To better understand the dynamics of this outbreak, 182 full genome HPAI H5N2 sequences isolated from commercial layer chicken and turkey production premises were analyzed using evolutionary models modified to incorporate epidemiological and geographic information. Epidemiological compartmental models constructed in a phylogenetic framework provided evidence that poultry type acted as a barrier to the transmission of virus among midwestern poultry farms. Furthermore, after initial introduction, a continuous external source of virus was not needed to explain the propagation of HPAI cases within the commercial poultry industries. Discrete trait diffusion models indicated that within state viral transitions occurred more frequently than inter-state transitions. Distance, road density and proportion of water coverage were all supported as associated with viral transition between county groups (Bayes Factor > 3.0). Together these findings indicate that the midwestern poultry industries were not a single homogenous population, but rather, the outbreak was shaped by poultry sectors and geographic factors.

**AUTHOR SUMMARY:** The highly pathogenic avian influenza outbreak among poultry farms in the midwestern United States appears to be influenced by agricultural and geographic factors. After initial introduction of the virus into the poultry industries, no further introductions (such as from a wild bird reservoir) were necessary to explain the continuation of the outbreak from March to June 2015. Additionally, evidence suggests that proximity increases the chances of viral movement between two locations. While many hypotheses have been proposed to explain the transmission of virus among poultry farms, the support for road density as an important driver of viral movement suggests human-mediated viral transportation played a key role in the spread of the highly pathogenic H5N2 outbreak in North America.

## INTRODUCTION

In 2014, a novel reassortant highly pathogenic avian influenza (HPAI) H5N8 virus of the hemagglutinin (HA) clade 2.3.4.4 was identified in South Korean poultry and wild birds and quickly spread to other Asian countries and Europe (1–3). By the end of 2014, both the Eurasian H5N8 virus and its reassortant H5N2 containing Eurasian- and North American-origin gene segments, were reported in western Canada and the United States (4–6). The ensuing 2014-2015 North American HPAI outbreak marked the largest and most expensive HPAI outbreak in the United States to date (7). In late November 2014, commercial poultry flocks in British Columbia, Canada were reported to be infected with the novel reassortant HPAI H5N2 (5), soon followed by HPAI H5N8 isolation within wild birds in the United States Pacific Northwest (4). Over the next several months, sporadic infections arose in wild and domestic birds, including both commercial production and backyard poultry operations. In March 2015, a drastic increase of HPAI H5N2 cases was observed within domestic poultry in the Midwestern United States. By the end of the outbreak in June 2015, over 50.4 billion poultry died or were culled due to the outbreak, costing the US government over $850 million, the poultry industries an estimated $700 million to $1 billion and had a negative $3.3 billion impact on the economy (7–9).

Risk factors that explain the continued transmission of HPAI between domestic poultry facilities remain unclear. For instance, previous analyses have provided conflicting evidence as to the role of wild birds in the propagation of the outbreak within the midwestern poultry industries. Despite frequent reports of wild birds on the grounds and within barns of HPAI-positive turkey premises (10), a case-control study found no significant difference in exposure to wild birds between positive turkey premises and matched controls (11). Similarly, one phylodynamic analysis found no evidence of continued HPAI introductions into the Midwestern poultry industries (12), but other models have suggested multiple introductions (13,14). Geographic and environmental variables, such as human population, agricultural, climatological, and ecological measures, may help explain farm-to-farm transmission observed within the poultry industries. For example, proximity between midwestern poultry premises has been implicated as an important risk factor for HPAI infection (11,13). Although it has been suggested that poultry production type did not affect outbreak transmission (12), this has not been formally tested. Despite extensive molecular epidemiological studies, such environmental and ecological covariates of viral spread during this outbreak have not been investigated.

Direct epidemiological links between most poultry premises have not been established (15), limiting the ability to investigate risk factors that facilitated HPAI transmission among poultry farms. The incorporation of pathogen genetic sequence data into epidemiological investigations can elucidate network connections between infectious entities, be that individual hosts or populations, such as poultry farms. One approach is viral phylodynamic modeling, ie. the integration of epidemiological and evolutionary models to explore viral ecological dynamics. Based on the assumption that viral epidemiology and evolution occur on the same time scale, viral phylodynamic modeling can reveal underlying population structure and epidemiological parameters. Recent incorporation of generalized linear models (GLM), a family of commonly used regression methods, into Bayesian phylogenetic frameworks have enabled investigation into the impact of ecological factors on the geographic diffusion of viral pathogens (16,17). Through such an approach, factors associated with HPAI movement within United States poultry industries can be identified, informing future control efforts.

In this study, we integrated epidemiological and ecological parameters with genomic sequence data collected contemporaneously with the midwestern poultry industries HPAI outbreak to formally test outstanding hypotheses. Whole genome HPAI H5N2 sequence data isolated from layer chicken and turkey premises were analyzed using evolutionary model-based techniques. First, we developed population models to test the importance of poultry sector divisions (i.e. layer chicken vs turkey industries) and external viral introductions from an unsampled avian population in the propagation of the outbreak. Second, we evaluated ecological predictors of geographic diffusion of virus among midwestern counties to help identify environmental and human variables associated with viral transmission. Together, these analyses use information that accumulated within the HPAI H5N2 genome during the outbreak to help decipher higher-order patterns of viral dispersal among commercial poultry farms.

## RESULTS

### HPAI H5N2 Evolution within Domestic Poultry

182 full genome HPAI H5N2 genetic sequences, each representing a single commercial poultry farm operation across 49 counties in six states (Iowa, Minnesota, Nebraska, North Dakota, South Dakota, and Wisconsin), were included in the present analysis. The sequences were isolated from samples collected between March 25 and June 15, 2015 from positive turkey premises (72.5%) and layer chicken farms (27.5%). Two molecular clock assumptions and three “traditional” coalescent models (i.e., constant population, exponential growth, and extended Bayesian skyline plot [EBSP]) were compared with marginal likelihood estimation (MLE) to evaluate the underlying population and evolutionary dynamics of the 2015 HPAI outbreak. The highly flexible EBSP coalescent with a strict molecular clock assumption had the best fit for the included sequence data (log(MLE) = −25884.06). The supported molecular clock assumption varied depending on the coalescent model employed. Statistical support between the relaxed and strict molecular clock assumptions was ambivalent for the constant and exponential coalescent models (log Bayes factor of relaxed compared to strict molecular clock (logBF_R-S_) = 0.06 and - 0.3, respectively; Fig 1A). Stronger evidence for the strict clock was observed when the EBSP coalescent was implemented (logBF_R-S_ = −4.45). Similarity between molecular clocks was also demonstrated by the limited impact of the molecular clock assumptions on phylogenetic tree parameter estimates such as evolutionary rate and time to the most recent common ancestor (TMRCA; Fig 1B and 1C). For example, under the EBSP coalescent, the relaxed mean clock rate was 6.84×10^−3^ substitutions per site per year (95% highest posterior density (HPD): 6.09×10^−3^ − 7.59×10^−3^) compared to the strict clock rate estimate of 6.77×10^−3^ substitutions per site per year (95% HPD: 5.98×10^−3^ − 7.56×10^−3^). In contrast, selection of the coalescent model influenced the TMRCA of the included sequences. Under the strict molecular clock, EBSP coalescent models estimated the TMRCA as March 1, 2015 (95% HPD: February 16 to March 10, 2015) while the remaining traditional coalescent models had a TMRCA at least two weeks earlier (Fig 1C).

**Fig 1.**
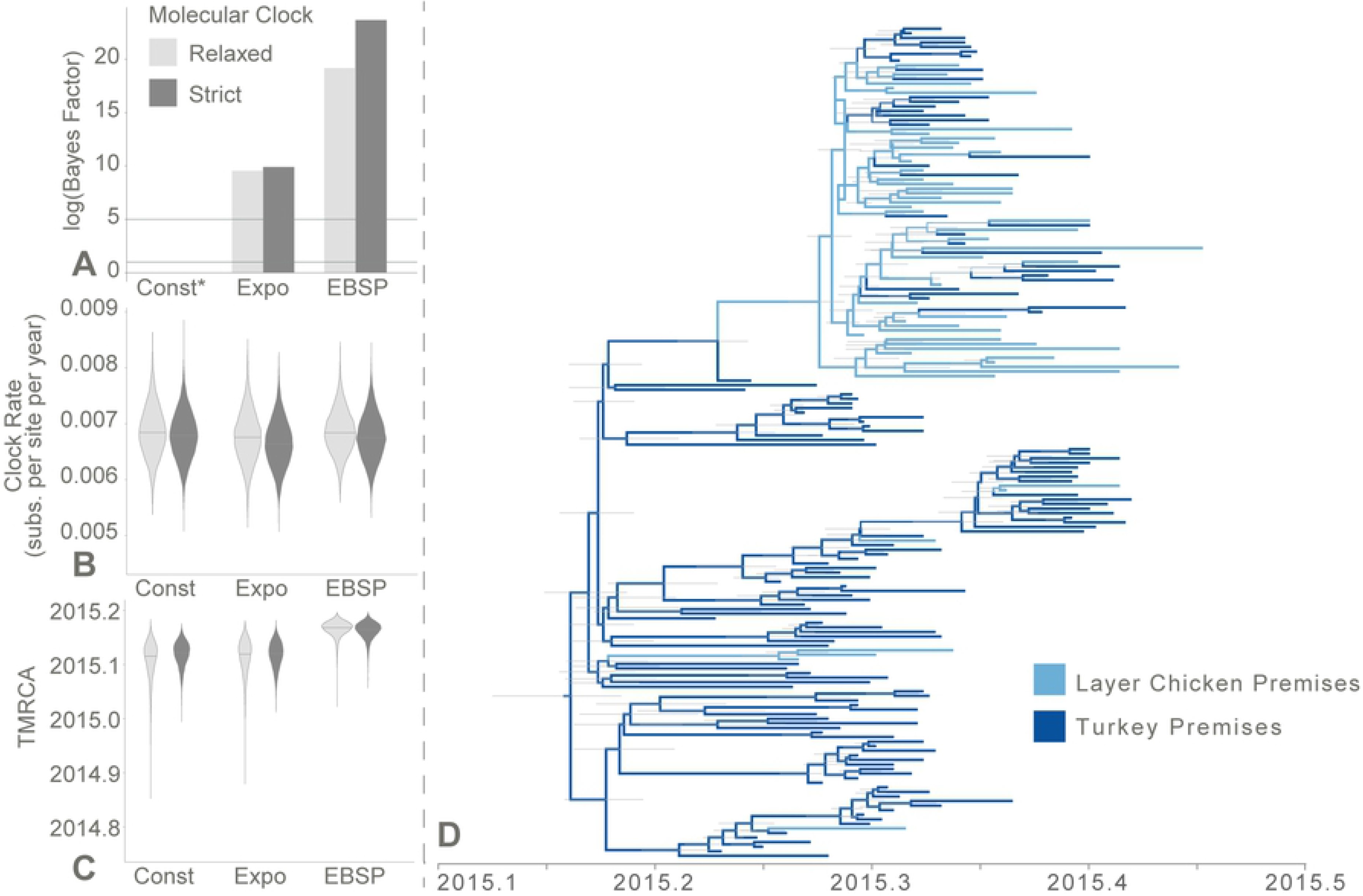
Evolutionary history of HPAI H5N2 isolated from commercial poultry premises, 2015. (A) Bayes factor (BF) tests between molecular clock and coalescent evolutionary models. For each coalescent model (exponential growth [Expo] and extended Bayesian skyline plot [EBSP]), BF was calculated using the constant coalescent model as reference (Const, indicated with asterisk) under the same molecular clock model. Two horizontal gray reference lines denote log(BF) = 1 and log(BF) = 5, which represent support and very strong support, respectively, for improved fit over the reference. (B) Molecular clock rate (substitutions per site per year) comparison between molecular clock and coalescent evolutionary models. (C) The estimated time of the most recent common ancestor (TMRCA; decimal year) compared between molecular clock and coalescent evolutionary models. (D) Maximum clade credibility tree representing the ancestral reconstruction of poultry industry (layer chicken vs. turkey) across the evolutionary history of the outbreak. The ancestral reconstruction assumed an EBSP coalescent and strict molecular clock evolutionary model. Tree branches are colored based on the most probable poultry industry of the descendant node. Thin gray node bars represent the 95% highest posterior density (HPD) of the node height (i.e., the time at which that ancestor is estimated to have existed).

### HPAI H5N2 Host Dispersion and Population Dynamics

To explore the extent of viral dispersal between poultry industries, multiple phylogenetic-based methods were performed: the structured coalescent, the discrete trait diffusion model, and epidemiologic compartmental model-based coalescent. Each of these methods estimate a different approximation for the dispersal of virus between populations. The structured coalescent treats layer chicken premises and turkey premises as separate population demes between which virus was allowed to “migrate,” and thus estimates a *migration rate* between the two demes. In contrast, discrete trait diffusion models treat the trait of interest (here, poultry industry) as a characteristic that evolves over time, inferring a *transition rate*, analogous to a nucleotide substitution model. Finally, compartmental models enable the calculation of *transmission rates* between the two poultry compartments. Although all approximate the amount of viral dispersal among the poultry industries, each measure is calculated differently with unique assumptions and so are referred to by a particular term. All methods estimated that viral dispersal from layer chicken premises to turkey premises occurred more frequently than from turkey premises to layer chicken (Supplemental Table S1). In the structured coalescent, the migration rate from layer chicken to turkey premises was much greater than the reverse (migration rate from chickens to turkeys: 12.6, 95% HPD: 6.2 – 18.7; migration rate from turkeys to chickens: 0.7, 95% HPD: 0.00001 – 2.2). The transition rates between the poultry industries estimated from the discrete trait diffusion model were much more similar to each other (transition rate from chickens to turkeys: 1.4, 95% HPD: 0.04 – 3.9; transition rate from turkeys to chickens: 0.3, 95% HPD: 0.003 – 0.9). These models suggest the dispersion of virus between poultry industries was not symmetrical, potentially indicating poultry type played a role in the outbreak dynamics.

To formally test this hypothesis, we used epidemiological compartmental model equations to describe the coalescent process (18). Four competing scenarios were constructed (Fig 2A). Models 1 and 2 described a homogenous poultry population that differed by the presence of a continuous external viral source in Model 2. In contrast, Models 3 and 4 described a host population stratified by poultry production system, again differing based on an external viral source in Model 4. It should be noted that due to the sampling scheme of genetic sequences (one HPAI whole genome sequence per infected premises), the epidemiologic unit of interest was the premises (or farm), and not the individual bird. That is, findings of the compartmental models should be interpreted on the farm-to-farm scale, not the dynamics of transmission between individual birds. Akaike’s information criteria for Markov chain Monte Carlo (AICM) calculated from the posterior sample of structured tree likelihood estimates revealed that Model 3 provided the best fit for the data under both strict and relaxed molecular clock assumptions (AICM under strict clock = 330.1; under relaxed clock = 376.3; Fig 2B, Supplemental Table S2). This suggests the midwestern portion of the 2015 HPAI outbreak was isolated from external sources but most likely structured by poultry production system. Four transmission rates were estimated for Model 3 to describe the interaction between the layer chicken and turkey populations: two within-poultry system rates (β_T_ and β_C_) and two between-poultry system rates (β_TC_ and β_CT_). The model estimated the transmission rates within the turkey production system to be highest (β_T_ = 11.6, 95%HPD: 2.0 – 22.0), followed by transmission rates from chicken farms to turkey farms (β_CT_ = 4.9, 95% HPD: 0.6 – 9.6). The lowest transmission rate was estimated from turkey farms to chicken farms (β_TC_ = 0.1, 95% HPD: 0.02 – 0.22). This is similar to the results of the structured coalescent model and discrete trait model described above (Supplemental Table S1). Infectious period of a farm also varied substantially between the two production systems. A HPAI-positive turkey premises was estimated to remain infectious for 5.7 days (95% HPD: 4.3 – 10.5), whereas layer chicken premises were estimated to remain infectious for 32.1 days (95% HPD: 22.4 – 49.3; Fig 2C).

**Fig 2.**
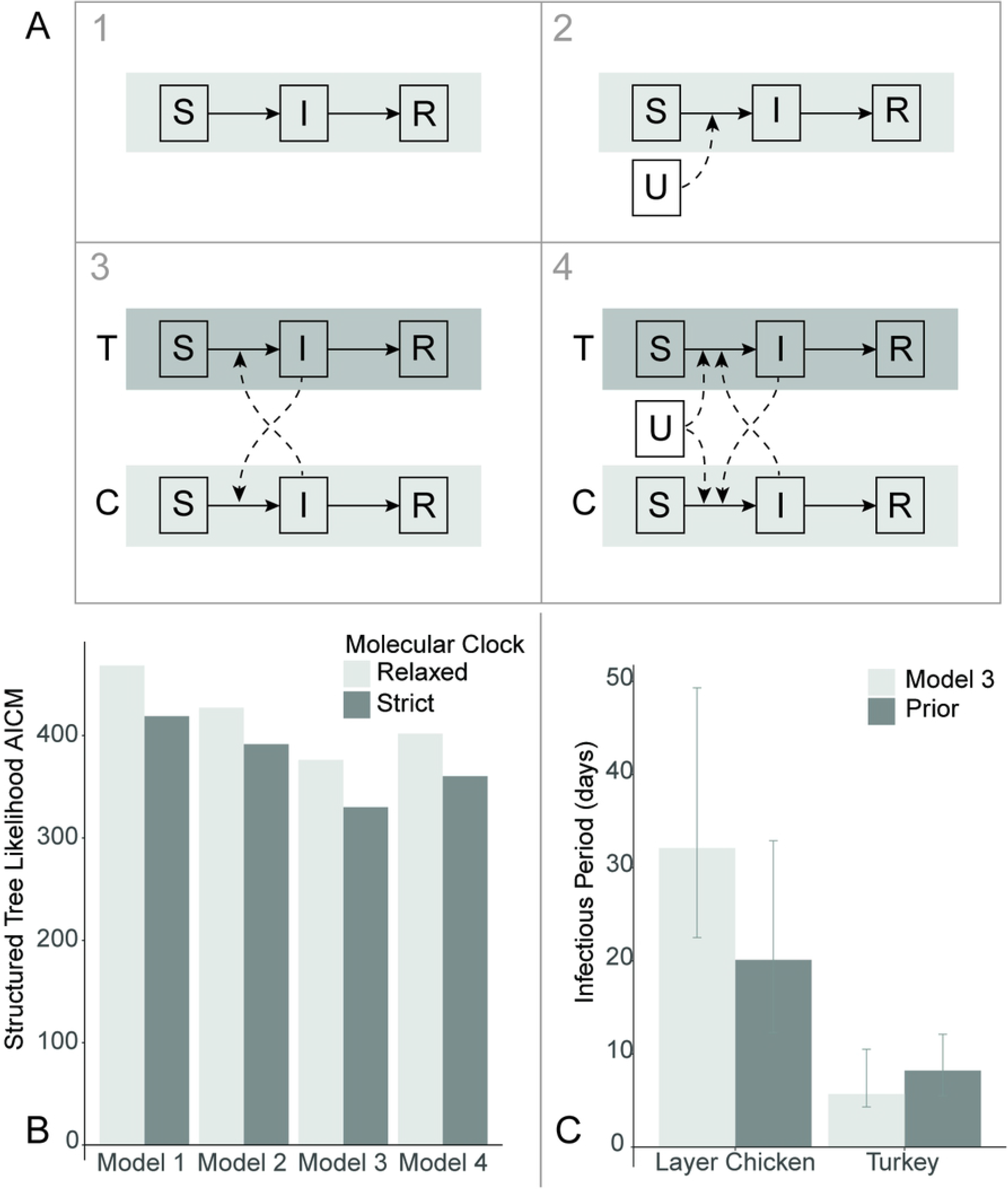
Comparison of hypothesized HPAI H5N2 epidemiological compartmental models. (A) Each compartmental model represents a Susceptible-Infectious-Removed (SIR) model with varied population heterogeneity: 1) a single, closed, homogenous population, 2) a single, homogenous population with a continual external source of virus (U), 3) a closed population, stratified by poultry system (turkeys (T) and layer chickens (C)), and 4) the stratified population with a continual external source of virus. (B) Compartmental model fit for the midwestern highly pathogenic avian influenza (HPAI) H5N2 outbreak, 2015. Akaike’s information criteria for Markov chain Monte Carlo (AICM) calculated based on the posterior distribution of the structured tree likelihood was used to evaluate the relative model fit for the four assessed compartmental models under differing molecular clock assumptions. Under both molecular clocks, Model 3 provided the best model fit. (C) Estimated infectious period of layer chicken and turkey farms during the 2015 midwestern highly pathogenic avian influenza (HPAI) H5N2 outbreak. During model specification, an informative prior was provided for the Bayesian process. This prior probability distribution was based on the reported average time from HPAI confirmation to depopulation plus 5 days to allow for delay between infection and HPAI confirmation. Model 3 estimated the infectious period for layer chickens to be longer than expected given the prior information.

### Ecologic Predictors of HPAI H5N2 Geographic Diffusion

Using the posterior distribution of phylogenetic trees estimated under the EBSP coalescent and strict molecular clock assumptions, discrete trait diffusion models were estimated to describe the geographic dispersal of HPAI H5N2 throughout the midwestern United States. County of origin was used as the basis to categorize the 182 sequences. Counties were grouped based on their state and whether sequences within the county exclusively originated from commercial turkey premises. For example, Iowan counties with only turkey cases were grouped separately from Iowan counties which had at least one layer chicken case. County groups with only turkey cases are henceforth referred to as turkey-exclusive while county groups with at least one layer chicken case are referred to as mixed poultry. The complete ancestral reconstruction of the midwestern outbreak is shown in Fig 3A. The three largest transition rates were observed between county groups within the same state, particularly Minnesota and Iowa (Fig 3B; Supplemental Table S3). The most frequent transitions occurred from Minnesota mixed poultry counties to Minnesota turkey-exclusive counties (median rate: 3.3 transitions per year; 95% HPD 0.7 – 6.4; BF = 490.6). In Minnesota, the reverse rate (i.e., from turkey-exclusive counties to mixed poultry counties) was also decisively supported with a relatively high transition rate (2.3 transitions per year; 95% HPD: 0.6 – 4.6; BF = 2,007.1). The second most frequent transition had the highest statistical support and occurred from Iowan mixed poultry counties to Iowan turkey-exclusive counties (3.3 transitions per year; 95% HPD 1.4 – 5.7; BF = 28,139.6). Three inter-state transitions were also decisively supported, but less frequent. These transitions were estimated from Iowa mixed-poultry counties to Minnesota turkey-exclusive counties (0.9 transitions per year; 95% HPD 0.2 – 2.2; BF = 14,068.3), from Minnesota turkey-exclusive counties to Wisconsin turkey-exclusive counties (0.74 transitions per year; 95% HPD 0.1 – 1.8; BF = 202.3) and from Wisconsin turkey-exclusive counties to Iowan mixed-poultry counties (0.9 transitions per year; 95% HPD 0.01 – 2.6; BF = 134.8). All supported transition rates (BF > 3.0) were found either within a state or between states that share borders, except for a single weakly supported rate from South Dakota turkey counties to Wisconsin mixed poultry counties (0.6 transitions per year; 95% HPD 0.0002 – 2.1; BF = 3.2). This suggests geographic distance influences the dispersal of HPAI H5N2 among midwestern counties.

**Fig 3.**
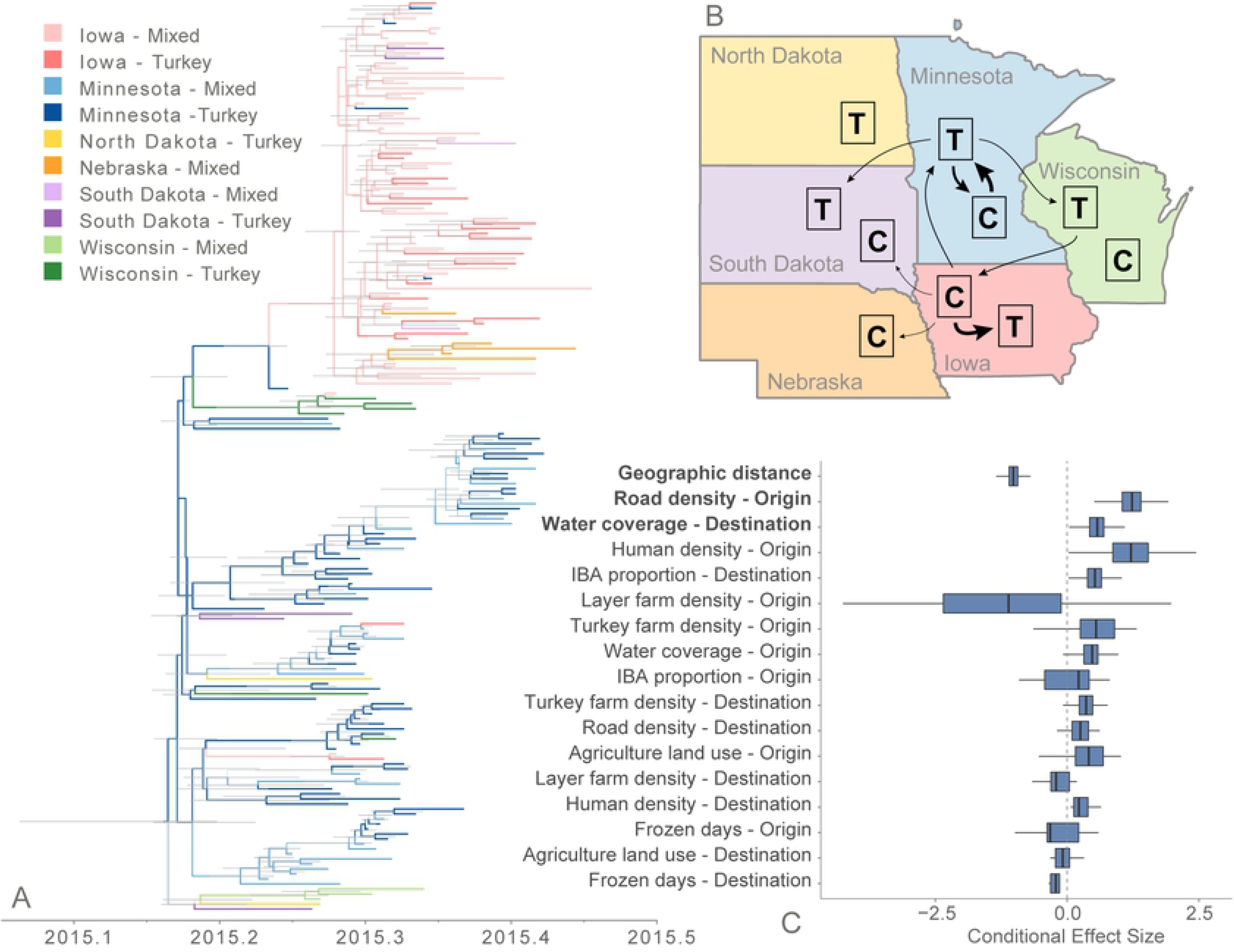
Discrete trait diffusion model of HPAI H5N2 among midwestern county groups. (A) Maximum clade credibility tree representing the ancestral reconstruction of county groups across the evolutionary history of the outbreak. The ancestral reconstruction was based on an EBSP coalescent and strict molecular clock evolutionary model. Tree branches are colored based on the most probable county group of the descendant node. Thin gray node bars represent the 95% highest posterior density (HPD) of the node height (i.e., the time at which that ancestor is estimated to have existed). (B) Diffusion rate summary among county groups. County groups were defined based on state and composition of host type within the county. Counties with only turkey cases (turkey exclusive; T) were grouped separately from counties with at least one layer chicken case (mixed poultry; C). Arrows represent transition rates with strong support (Bayes factor > 10) with arrow thickness proportional to the magnitude of transition rate. (C) Conditional effect size of environmental and geographic covariates within the generalized linear model (GLM). Conditional effect size represents the effect size of the variable coefficient given inclusion in the GLM. Supported covariates (Bayes factor > 3) are bolded. Covariates are ordered by Bayes factor. The dashed gray line represents a conditional effect size of 0, signifying little impact of the covariate on viral dispersal.

The discrete trait diffusion model was extended with a GLM that assessed the impact of distance and other environmental variables on the transition rates among the defined county groups. County characteristics for the 9 modeled variables are summarized in Table 1. On average, county centers were 266 km apart, ranging from 30 to 862 km. HPAI-positive counties had a higher density of layer chicken farms (0.02 farms/km^2^) than turkey farms (0.004 farms/km^2^). Counties also had a broad range of human population density ranging from about 1 to 58 people/km^2^. Of the 9 variables included in the GLM, three were statistically supported to be associated with diffusion of HPAI H5N2 among county groups (Fig 3C, Supplemental Table S4). Distance between county group centroid was decisively supported to be negatively associated with transition between two groups (log conditional effect size = −1.0; 95% HPD −1.2, −0.8; BF = 216,262.9). In other words, viral transitions are less likely between county groups that are separated by a greater distance. Road density of the origin county group was positively associated with viral dispersion (log conditional effect size = 1.2; 95% HPD 0.6 – 1.7; BF = 42.8). That is, county groups with a higher density of roads were associated with higher dispersion rates to other county groups. The proportion of the destination county group covered with water was only weakly supported for inclusion in the GLM (log conditional effect size = 0.6; 95% HPD 0.2 – 0.9; BF = 3.9).

**Table 1.** Demographic and geographic characteristics of the 49 United States counties with HPAI-positive commercial poultry premises during the H5N2 outbreak, 2015.

|  | **Mean** | **Standard**<br>**Deviation** | **Minimum** | **Maximum** |
| --- | --- | --- | --- | --- |
| **Distance between counties (km)** | 265.95 | 153.45 | 30.18 | 861.99 |
| **Layer Chicken Farm Density (farms/km<sup>2</sup>)** | 0.02 | 0.02 | 0.001 | 0.09 |
| **Turkey Farm Density (farms/km<sup>2</sup>)** | 0.004 | 0.004 | 3.97x10<sup>-6</sup> | 0.01 |
| **Human Population Density (humans/km<sup>2</sup>)** | 12.53 | 11.21 | 1.39 | 58.07 |
| **Road Density (km/km<sup>2</sup>)** | 1.89 | 0.32 | 1.18 | 2.78 |
| **Water Coverage (%)** | 1.56 | 2.21 | 0.02 | 11.34 |
| **Important Bird Area (%)** | 4.23 | 7.60 | 0.0 | 30.96 |
| **Agricultural Land Use (%)** | 78.07 | 13.28 | 32.46 | 90.62 |
| **Frozen days** | 19.82 | 4.93 | 12 | 38 |

## DISCUSSION

Our exploration of population models to describe the 2015 midwestern United States HPAI H5N2 outbreak provides evidence that upon entering the midwestern poultry industries, no further viral introductions from outside sources were needed to explain the observed epidemiological trajectory. Furthermore, the statistical support for a stratified poultry population suggests that poultry industries should not be considered a homogenous host population for viral pathogens. This is also supported by the discrete trait diffusion analyses, which demonstrate that geographic factors influence viral dispersion among counties, indicating heterogeneity among geographic locations. Multiple factors including poultry production system barriers and geographic characteristics appear to have influenced the course of the outbreak within poultry industries.

In our analysis, the EBSP coalescent model had better support than the other traditional coalescent models in terms of model fit. This is most likely a reflection of EBSP’s flexibility, i.e. the piece-wise nature of this method, which facilitates the identification of complex population changes. Coalescent theory has been a popular technique to infer population demographics underlying viral outbreaks (19–28). By relating effective population size to the rate at which phylogenetic lineages converge backwards in time, the coalescent has become a powerful tool to infer demographic changes even in the face of incomplete sampling. Traditionally, the estimation of the coalescent process required rigid prior assumptions in the form of simplistic mathematical growth functions (e.g., constant population size or exponential growth). To better reflect biological reality, methods have been developed that incorporate more flexibility than a one to two parameter mathematical function (29,30). For instance, the EBSP assumes demographic changes follow a smoothed, piece-wise, linear function whose change points are inferred from the sequence data (31). To date, mathematical methods to incorporate population structure into EBSP coalescent models have not been developed even though population structure has been observed to confound EBSP estimates (32).

Despite the flexibility of EBSP, compartmental-based coalescent models are worth assessing as they allow for direct incorporation and hypothesis testing of specific population structures. Rather than the non-parametric, piece-wise approach of EBSP, the prior mathematical functions assumed are ordinary differential equations (ODEs) constructed from the specification of epidemiological compartmental models. It is the parameters of these ODEs that are fit during the Markov chain Monte Carlo (MCMC) process. Among the four analyzed compartmental models, we found that the closed, stratified population provided the best fit for the sequence data, suggesting layer chickens and turkeys represented two separate host populations that interacted with each other, but did not receive virus from a continuous external source. Interestingly, when only observing the single homogenous population models (Models 1 and 2), the inclusion of an external viral source (Model 2) improves model fit compared to the closed population model (Model 1). Once the population structure of poultry type is included (Models 3 and 4), the closed population model provides a better fit than that with continual viral introductions. This observation underlines the importance of including population heterogeneity within evolutionary demographic models to explain observed viral diversity and population dynamics.

To help improve identifiability of the remaining parameters within the compartmental model, expected prior distributions for the infectious period of affected premises were specified based on reported USDA data (7). Despite the informative assumption, the infectious period of layer chicken farms was estimated to be longer than expected. In our model, we assumed a 5-day period between the onset of infectivity of the farm and reporting of HPAI infection. Delays in the identification and/or reporting of HPAI infection could result in infectious periods that begin well before the assumed 5 days. Continued infectivity beyond the completion of flock depopulation is another likely contributor to prolonged infectious periods. Although commercial poultry depopulation occurred on average 6.4 days after National Veterinary Services Laboratory (NVSL) HPAI confirmation, premises were not considered to be virus-free until, on average, 87.7 days following confirmation (7). In either case, our models suggest layer chicken farms remained infectious for much longer than turkey farms, potentially explaining why the transmission rate from chicken farms to turkey farms was higher than its counterpart. In fact, regardless of the model (i.e., structured coalescent, discrete trait diffusion model, or compartmental model), layer chicken farms played a more central role to viral transmission than turkey farms during the outbreak. This may seem contradictory to experimental evidence that demonstrated the HPAI H5N2 virus had longer mean death times in turkeys (5 – 6 days) compared to chickens (2 – 3 days (33). However, such experimental infections only describe transmission information on the individual bird scale, rather than the farm-to-farm transmission scale captured in this analysis. Although it may be that individual turkeys survive longer, in practice turkey premises were quicker to be depopulated, resulting in a shorter farm-level infectious period compared to chicken farms. Because intervention (i.e. depopulation) was performed on the farm level, individual-level infectious periods alone are not adequate to describe the overall observed outbreak dynamics.

The implementation of a GLM into a Bayesian discrete trait analysis has been previously applied to HPAI in China (34) and Egypt (35), providing evidence that environmental, agricultural and anthropogenic factors influence viral movement. Due to differences in social, governmental and agricultural systems, the generalization of these previous GLM results to other countries may not be appropriate. Instead, these studies provide a framework to identify epidemiological covariates of the North American HPAI H5N2 outbreak. Our results indicate that distance and road density are key factors that influenced the geographic spread of HPAI H5N2 among midwestern counties in the spring of 2015. A recent spatial modeling analysis revealed that HPAI spread among Minnesota poultry premises was heavily distance-dependent during the 2015 outbreak (13). Our results support this claim by providing evidence that the frequency of shared viral diversity increases as the distance between two counties decreases. Risk of infection due to proximity can also be observed in our discrete trait diffusion model in which within-state HPAI spread was much more frequent than inter-state spread. HPAI movement between states may explain Bonney, et al.’s finding that distance-independent transmissions significantly improved the fit of their transmission kernel model (13). Although the causal relationship between the supported covariates and viral dispersal cannot be determined from our analysis, the statistical support for road density within the GLM may provide evidence for the relative importance of anthropogenic movement of virus. High road density may correlate with better logistic connectivity between farms, increasing the likelihood that an infected premises will export virus to nearby farms and counties. Road density has been associated with HPAI H5N1 outbreaks in Bangladesh (36), Thailand (37), Romania (38), Indonesia (39,40), and Nigeria (41), although high road density in these countries may reflect greater human population density and, therefore, a higher likelihood of case detection (42). Intensive commercial poultry surveillance during the 2015 outbreak and the lack of support for human population density as a covariate within our model suggest that the statistical support for road density in the dispersal of HPAI among midwestern counties may not merely be an artifact of sampling bias or confounding. The third variable associated with HPAI dispersal was the proportion of a destination county group covered by surface water. In other words, counties with a larger proportion of surface water received virus more frequently compared to those with less surface water. Surface water resources have been associated with HPAI dispersal and prevalence in China and may signify movement of virus by migrating waterfowl that stopover in lakes, rivers and wetlands (34,43). In our analysis, this variable was only weakly supported and had a relatively small effect size. Additionally, other variables that represent potential migratory stopover habitats, such as Important Bird Areas and agricultural land, were not supported within the model, suggesting that if wild birds contributed to HPAI dispersal within the Midwest, their role was limited. This further supports previous studies, which indicated that the Midwestern portion of the outbreak was driven by inter-farm transmission (11,12,14). Several mechanisms have been proposed to explain HPAI transmission between farms during the 2015 outbreak, including equipment sharing, personnel overlap, and aerosolization.

Due to the restricted number of sequences in the presented analysis, the number of variables and demographic scenarios that could be modelled was limited. This also affected the resolution of the geographic covariates that could be included within the GLM. Ideally, the environmental and agricultural characteristics of each individual farm or county would be evaluated as predictor for HPAI spread; however, the individual transition rates between 182 farms or even 49 counties would be impossible to accurately estimate from the 182 sequences of this dataset. For this reason, sequences were categorized into county groups, resulting in a manageable transition rate matrix as well as permitting the summarization of environmental characteristics across a few counties rather than across an entire state.

Despite these limitations, our results present several implications for future HPAI surveillance and control in the United States. While wild birds may provide a means of viral dispersal across large distances and initial introduction into an area, evidence suggests the HPAI outbreak within the midwestern poultry industries could be maintained without continued introductions. In this sense, in-place biosecurity efforts may have been enough to prevent continued viral introductions from outside sources (including wild birds, backyard poultry flocks, or long-distance movement from other geographic regions), but were ineffective against local farm-to-farm transmission. For instance, it has been suggested that biosecurity factors could explain the lack of HPAI cases within the broiler chicken industry in the Midwest (44). The association of distance between and road density of county groups with HPAI dispersal suggests human transportation modes may have played an important role in dispersal of HPAI across the Midwest. A better understanding of how HPAI-positive farms are logistically connected would greatly aid surveillance and control efforts. With the knowledge of how these farms share personnel and equipment, future outbreaks could be contained by disruption of the transportation network.

## METHODS

### Dataset

Whole genome HPAI H5N2 sequences collected, isolated and sequenced by the United States Department of Agriculture (USDA) during the 2014 – 2015 North American HPAI outbreak served as the basis for the analyzed data set. Full description of their collection and sequencing has been reported elsewhere (14). A subset of this sequence data was selected to better investigate the farm-to-farm transmission dynamics of the midwestern portion of the HPAI H5N2 outbreak. This subset was defined by the following inclusion criteria: 1) sequences isolated from commercial domestic poultry samples and 2) membership of the sequence in a phylogenetically distinct group, as determined by maximum likelihood estimation by Lee, et al (14). These viruses represented midwestern HPAI-positive poultry premises from the latter part of the outbreak, which was defined by a rapid increase in incidence within the midwestern poultry industries. As within-farm epidemiological dynamics were not of interest in this analysis, only one viral sequence per positive poultry premises was included. Viruses isolated from backyard poultry operations and wild birds were not included due to the incongruency in surveillance and sampling between these populations and the domestic poultry industries. A full list of the included sequence names and accession numbers are provided in Supplemental Table S5.

### Coalescent Model Comparison

Coalescent theory provides the statistical framework to estimate population changes over time from genetic sequence data. To investigate the population dynamics of the midwestern poultry portion of the outbreak, various coalescent population model prior assumptions were implemented and compared in BEAST2 (45). Using ModelFinder (46) as implemented in the IQ-TREE software package (http://www.iqtree.org/), the Kimura three parameter (K3P; i.e., one transition rate and 2 transversion rates) model (47) with unequal base frequencies and a gamma distribution of rate variation among sites (48) was determined as the best fit nucleotide substitution model and was used for each BEAST2 model. All coalescent models were separately estimated under both strict and lognormally distributed, uncorrelated, relaxed molecular clock assumptions. For each BEAST2 model, at least three independent MCMC runs of 50 million chain length were initiated from random starting trees. Convergence was assessed in Tracer v1.5, ensuring an effective sample size (ESS) > 200 for each estimated parameter. If ESS < 200, the discarded burn-in fraction was increased or more MCMC runs were performed. Three “traditional” coalescent models (i.e., constant population, exponential growth, and EBSP (31)) were performed to investigate demographic dynamics. Model fit was compared among the coalescent and molecular clock models with path sampling to calculate the marginal likelihood estimate (MLE) (49). Estimating the marginal likelihood enables the calculation of a Bayes Factor (BF), which is a ratio of two marginal likelihoods. A log(BF) > 5 indicates very strong statistical support for one model over the other (50). Viral dispersion between poultry industries (layer chicken vs. turkey) was initially estimated with a simple discrete trait diffusion model as well as a structured coalescent (25). The EBSP coalescent model was used as the tree prior for the discrete trait diffusion model. Both viral dispersion models were performed under both strict and relaxed molecular clock, as above.

A recently developed structured coalescent-based BEAST2 package (PhyDyn) was used to investigate more complex pathogen population scenarios by specifying epidemiological compartmental models (18). Four alternative compartmental models were assessed to investigate the presence of population structure by poultry type (layer chicken vs. turkey) and continual viral introductions from an unknown source population. Each compartmental model was a Susceptible-Infectious-Removed (SIR) model with varied population heterogeneity (Fig 2A): 1) a single, closed, homogenous population, 2) a closed population, stratified by poultry system, 3) a single, homogenous population with a continual external source of virus, and 4) a stratified population with a continual external source of virus. By including models with an external viral source, the models test whether this aspect of the outbreak was insulated or involved repeated introductions of HPAI from wild birds, backyard poultry, or undetected HPAI-positive premises. Since marginal likelihood estimation via path sampling has not yet been developed for the PhyDyn package, Akaike Information Criterion for MCMC (AICM) (51) was used to assess model fit and was calculated from the posterior MCMC sample of the structured tree likelihood with the R package, aicm (https://rdrr.io/cran/geiger/man/aicm.html).

### Discrete trait diffusion models

To estimate the impact of environmental variables on the geographic diffusion of HPAI between midwestern counties, a discrete trait diffusion model was constructed and further extended with a generalized linear model (GLM) in BEAST v1.10 (52). Discrete trait diffusion models are a phylogeographic technique in which each analyzed genetic sequence is assigned an observed characteristic trait that is assumed to have changed across the viral evolutionary history in a continuous time Markov chain process (53). Transition rates among these observed traits can then be inferred. In this analysis, the discrete character trait definition was based on the United States county in which the HPAI-positive poultry premises was located. Counties were then categorized by state and whether the county’s sequences exclusively originated from turkey production premises. In contrast to the simplified discrete trait model performed parallel to the structured coalescent model above, this model enables geographic dispersion of the HPAI virus to be estimated.

The geographic discrete trait diffusion model was extended with a GLM to assess the impact of environmental covariates on the viral transition rates among county categories. In this approach, viral diffusion rates among discrete geographic regions act as the outcome to a log-linear combination of environmental variables, regression coefficients and indicator variables (17). Environmental and anthropogenic variables were selected based on previous indication of their importance to avian influenza risk (42). Layer chicken farm density and turkey farm density were calculated from USDA 2012 census data (https://quickstats.nass.usda.gov/) divided by the land area of the county group. Human population density and proportion of county covered in water was obtained from United States census data (https://factfinder.census.gov/). The remaining variables were summarized per county group using ArcGIS Pro. Geographic distance was calculated as the linear distance between county group centroid. Road density was estimated as the total length of road per county divided by the total county group area. Proportion of county designated as an important bird area (IBA) was calculated using the publicly available Audubon Important Bird Areas and Conservation Priorities data (54). Proportion of the county group used for agriculture (i.e., covered by pasture, hay or cultivated crops) was obtained from the United States Geological Survey National Land Cover Database created in 2011 and amended in 2014 (55). The number of frozen days was calculated from daily freeze-thaw satellite data from March 1 to June 15, 2015 (56,57). A frozen day was defined as a day in which more than half of the county group area had a temperature measured as below 0 C. All covariate measures were log-transformed and standardized before inclusion in the GLM.

The discrete trait diffusion models were applied to the empirical distribution of phylogenetic trees from the best fitting evolutionary model. For each diffusion model, three independent MCMC runs of 1 million steps in length were performed, sampling every 100 steps. Convergence was assessed in Tracer v1.5, ensuring ESS > 200 for each estimated parameter. Removing the first 10% of each run as burn-in and re-sampling every 300 steps, log and tree files were combined using LogCombiner in the BEAST v1.10 software suite. Statistical support for transition rates in the discrete trait diffusion model and the covariate coefficients of the GLM were inferred using Bayesian stochastic search variable selection (BSSVS). Briefly, for each estimated parameter, an indicator variable (I) is stochastically turned on (I = 1) or off (I = 0) at each step of the MCMC (16,53). The posterior distribution of indicator values can be used to calculate a Bayes factor (BF), indicating the level of statistical support for the inclusion of that parameter in the model. BF support was defined in the following categories: no support (BF < 3.0), substantial support (3.0 ≤ BF < 10.0), strong support (10.0 ≤ BF < 30.0), very strong support (30.0 ≤ BF < 100.0), and decisive support (BF ≥ 100.0). Median transition rates, median conditional coefficients, 95% highest posterior density (HPD) and BF were calculated using personalized Python scripts.

## ACKNOWLEDGEMENTS

We would like to thank Alex Heri for her critical review of this manuscript.

## SUPPLEMENTAL MATERIAL CAPTIONS

Table S1. Estimates of viral transmission between poultry industries during the 2015 highly pathogenic avian influenza virus H5N2 outbreak within the midwestern United States.

Table S2. Akaike’s information criteria for Markov chain Monte Carlo (AICM) for the epidemiological compartment-based coalescent models.

Table S3. Discrete trait diffusion matrix of the midwestern highly pathogenic avian influenza (HPAI) H5N2 outbreak, 2015. Median rates and associated 95% highest posterior density intervals (in brackets) are presented in each cell. The diffusion model is asymmetrical, and therefore, rates have directionality from a source county group (indicated on the left) to a sink county group (indicated across the top). County groups were defined by state (IA - Iowa, MN - Minnesota, ND - North Dakota, NE - Nebraska, SD - South Dakota, WI - Wisconsin) and composition of poultry type (T - turkey exclusive, CM - layer chicken exclusive and mixed poultry). Rates are colored by the level of Bayes factor support. Gray rates represent no support.

Table S4. Generalized linear model (GLM) conditional effect sizes and statistical support for agricultural and geographic covariates of the dispersal of highly pathogenic avian influenza (HPAI) H5N2 among midwestern county groups. Conditional effect size and 95% highest posterior density (HPD) were calculated based on the estimated GLM coefficients given the Bayesian stochastic search variable selection (BSSVS) indicator = 1. The posterior probability (PP) refers to the proportion of Markov chain Monte Carlo (MCMC) samples in which the BSSVS indicator = 1. Bayes factor (BF) > 3.0 indicates statistical support for the inclusion of the covariate within the GLM.

Table S5. Accession number and names of 182 included HPAI H5N2 full genome sequences.

